# Gait Abnormalities and Aberrant D2 Receptor Expression and Signaling in a Mouse Model of the Human Pathogenic Mutation *DRD2^I212F^*

**DOI:** 10.1101/2022.06.09.495548

**Authors:** Dayana Rodriguez-Contreras, Sheng Gong, Joseph J Lebowitz, Lev M Fedorov, Naeem Asad, Timothy M Dore, Christopher P Ford, John T Williams, Kim A Neve

## Abstract

A dopamine D2 receptor mutation was recently identified in a family with a novel hyperkinetic movement disorder (Mov Disord **36**: 729-739, 2021). That allelic variant D2-I^212^F is a constitutively active and G protein-biased receptor. We now describe mice engineered to carry the D2-I^212^F variant, *Drd2^I212F^*. The mice exhibited gait abnormalities resembling those in other mouse models of chorea and/or dystonia, and had decreased striatal D2 receptor expression. Electrically evoked IPSCs in midbrain dopamine neurons and striatum from *Drd2^I212F^* mice exhibited slow onset and decay compared to wild type mice. In the presence of dopamine, current decay initiated by photolytic release of sulpiride from CyHQ-sulpiride was slower in midbrain slices from *Drd2^I212F^* mice than *Drd2*^+/+^ mice. Furthermore, in contrast to wild type mice in which dopamine is more potent at neurons in the nucleus accumbens than in the dorsal striatum, reflecting activation of Gα_o_ vs. Gα_i1_, dopamine had similar potencies in those two brain regions of *Drd2^I212F^* mice. Repeated cocaine treatment, which decreases dopamine potency in the nucleus accumbens of wild type mice, had no effect on dopamine potency in *Drd2*^I212F^ mice. The results demonstrate the utility of this mouse model for investigating the role of pathogenic *DRD2* variants in early-onset hyperkinetic movement disorders.

## Introduction

Many movement disorders are treated with or caused by drugs that modulate the activation of one or more dopamine receptors (Luquin-Piudo and Sanz, 2011; Cepeda *et al.*, 2014; Vaiman *et al.*, 2021). Nevertheless, no naturally occurring mutation of a dopamine receptor was known to cause a movement disorder until the recent identification of the *DRD2* variant c.634A > T, p.Ile212Phe (Fig. 1A), which co-segregates with a phenotype of progressive chorea and cervical dystonia in a Dutch family (van der Weijden *et al.*, 2021a). The dopamine D2 receptor modulates both Gα_i/o_-mediated signaling pathways (e.g., inhibition of adenylyl cyclase and activation of G protein-regulated inwardly rectifying potassium channels (GIRKs)) and arrestin-mediated signaling pathways (e.g., the protein kinase Akt and glycogen synthase kinase-3) (Beaulieu and Gainetdinov, 2011). The mutant D2-I^212^F exhibits enhanced activation of G protein-mediated signaling combined with diminished binding of arrestin in human embryonic kidney (HEK) 293 cells; thus, it is a constitutively active and signaling-biased receptor relative to the reference D2 receptor (Rodriguez-Contreras *et al.*, 2021; van der Weijden *et al.*, 2021a).

**Figure 1.**
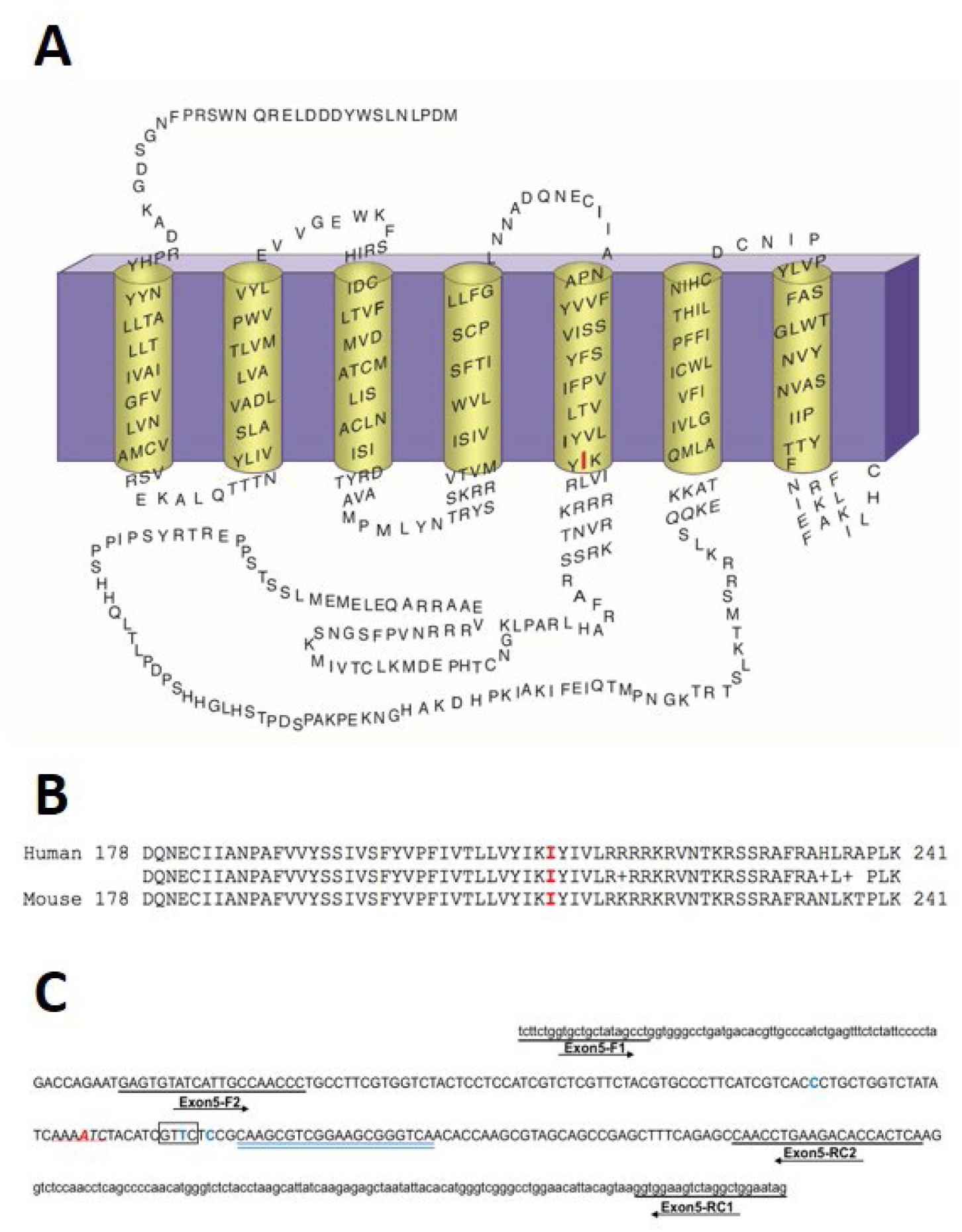
Strategy for Producing *Drd2^I212F^* mice. **A**, Membrane topology of the human D2L receptor, depicting the I^212^ position in red. **B**, Alignment of amino acid sequences from exon 5 of human and mouse D2 dopamine receptor genes, corresponding to the amino acids 178 to 241 in both proteins. Identity between human and mouse amino acid sequences is shown. **C**, Mouse *Drd2* exon 5 nucleotide sequence, including intron/exon 5 junctions, showing the PCR primers (underlined) used for genotyping the mice and the position of the gRNA sequence (double-underlined in blue). Exon 5 sequence is in uppercase and intron sequence in lowercase. In red is shown the position of the wild type amino acid (A-B) and nucleotide (C) that was changed to generate the D2-I^212^F variant. The single stranded oligonucleotide (ssODN) template contained three synonymous changes C-A, T-A and C-A; the WT nucleotide is shown at these positions in blue. Moreover, underlined sequence in red (C) shows the position of ApoI (RAATTY) site, whereas the box shows the location of the introduced RsaI (GTAC) site. Both restriction sites are present in KI but not in WT mice. R means purine (A/G) and Y, pyrimidine (T/C).

The enhanced D2-I^212^F-mediated activation of G proteins is manifested in both increased basal activation of Gα_i1_ and Gα_oA_, and increased agonist potency for activation of Gα_i1_ but not Gα_oA_ (Rodriguez-Contreras *et al.*, 2021). We speculated that this would affect striatal D2 receptor signaling. In striatal D2 receptor-expressing medium spiny neurons (D2-MSNs), the more efficient coupling of wild type D2 receptors to Gα_o_ in the nucleus accumbens (NAc) results in more potent dopamine activation of G protein-mediated signaling compared to that in dorsal neostriatum, which is mediated by Gα_i_ (Marcott *et al.*, 2018). Furthermore, repeated cocaine exposure decreases the abundance of Gα_o_ in the NAc and eliminates high-potency dopamine activation of GIRK that is mediated by wild type D2 receptors (Gong *et al.*, 2021). Because the high-potency coupling of D2-I^212^F to Gα_i1_ results in similar potency for activation of Gα_i1_ and Gα_oA_ in HEK 293 cells, we suggested that the potency of dopamine would not differ between the nucleus accumbens and dorsal striatum in mice expressing D2-I^212^F, and that accumbal dopamine signaling might be unaffected by repeated cocaine treatment (Rodriguez-Contreras *et al.*, 2021).

Gait abnormalities are common in movement disorders with choreatic and dystonic features, and sometimes treated with drugs that modulate dopamine receptor activity (Koller and Trimble, 1985; Barbosa and Warner, 2018). Gait abnormalities that have been observed in mouse models of dystonia and chorea include altered stride length and frequency (Dai *et al.*, 2009), the ratio of the time spent in propulsion to the time spent braking (Wright *et al.*, 2015), and splaying of the hind limbs (Liu *et al.*, 2015).

To evaluate the pathogenic role of the D2-I^212^F variant, we have now generated a knock-in mouse line, *Drd2^I212F^.* Both *Drd2^+/I212F^* and *Drd2^I212F/I212F^* mice exhibited gait abnormalities and decreased striatal D2 receptor expression. We also used this *Drd2^I212F^* mouse line to confirm and extend our prior characterization of the cellular effects of this pathogenic dopamine D2 receptor variant. GIRK-mediated inhibitory postsynaptic conductances (IPSCs) in *Drd2^I212F^* mice were characterized by slow kinetics in both midbrain dopamine neurons and D2-MSNs in the basal forebrain. In addition, cocaine treatment had no effect on the potency of dopamine in the basal forebrain MSNs of *Drd2^I212F^* mice, in contrast to cocaine effects in WT mice (Marcott *et al.*, 2018; Gong *et al.*, 2021), which is consistent with the higher potency of D2-I^212^F than D2-WT for Gα_i1_ observed in HEK293 cells (Rodriguez-Contreras *et al.*, 2021; van der Weijden *et al.*, 2021a).

## RESULTS

### Generation of *Drd2*^+/I212F^ mice

To mimic the human D2-I^212^F variant, we introduced the c.634A>T mutation in mice (Figure 1A) using the CRISPR-Cas9-mediated gene editing system (Jinek *et al.*, 2012). A detailed description of the strategy for producing the *Drd2^I212F^* knock-in (KI) mouse line, the genetic characterization of the mice, and confirmation of the probable absence of off-target effects can be found in Methods and Materials (see also Figure 1 – figure supplement1). We amplified and sequenced the exon 5 target locus (Figure 1B and C) from several heterologous *Drd2^+/I212F^* F0 mice and selected two founders (F0-429, lineage **A**, and F0-421, lineage **B**). Both founders, heterozygous for the mutation c.634A>T, were crossed with inbred control mice, and F1 pups were also genotyped by PCR/digestion and DNA sequencing (see details in Methods and Materials). *Drd2^I212F^* knock-in (KI) mouse lineages were maintained as heterozygous breeding colonies at the VA Portland Health Care System (VAPORHCS).

*Drd2^I212F/I212F^* and *Drd2^+/I212F^* mice (from both lineages) in their home cages were indistinguishable from *Drd2^+/+^* mice, showing no obvious defects in movement, size, or morphology. Of 179 mice born at the VAPORHCS of heterozygote crosses, the genotype distribution (*Drd2^+/+^*:*Drd2^+/I212F^*:*Drd2^I212F/I212F^*) was 32:51:30 for mice of lineage **A** and 16:35:15 for mice of lineage **B**, close to the expected 1:2:1 distribution.

### Gait Abnormality in *Drd2^I212F^* mice

In humans, the novel *DRD2^I212F^* variant co-segregates with a unique hyperkinetic movement disorder characterized by combined progressive chorea and cervical dystonia with adolescent onset. To quantify gait parameters in *Drd2^I212F^* mice, thirty-eight animals were tested on the DigiGait System, in which the mouse is recorded via a ventrally mounted camera while running on a transparent treadmill, at 10-12 months of age. The genotype distribution was 13 *Drd2^+/+^* (8 male and 5 female), 16 *Drd2^+/I212F^* mice (9 male and 7 female), and 9 *Drd2^I212F/I212F^* mice (4 male and 5 female).

Based on published results in other mouse models of chorea or dystonia (Dai *et al.*, 2009; Liu *et al.*, 2015; Wright *et al.*, 2015), we analyzed swing time, propel time, brake time, the ratio of time spent in propulsion to time braking, stride frequency, stride length, and stance width.

Results from left and right limbs were pooled (except for stance width), whereas fore- and hind limbs were analyzed separately. There was a significant effect of genotype (ANOVA) from 5 of the 7 measures for forelimbs and 3 for hind limbs, with increased stride length and decreased stride frequency being the most robust genotype-dependent effects observed (Table 1). For *Drd2^+/I212F^* mice, post hoc analysis identified significant increases in forelimb swing time, hind limb propel time, and forelimb and hind limb stride length, as well as decreased forelimb and hind limb stride frequency. Homozygous mutant *Drd2^I212F/I212F^* exhibited the same changes and also significantly increased forelimb propel time and propel/brake ratio (Table 1).

**Table 1.** Gait analysis in *Drd2^I212F^* mice. Results are shown for selected measures of gait obtained using a DigiGait treadmill. Time measurements (Swing, time spent swinging the limb forward; Propel, time spent in propulsion with the paw pushing backward; Brake) are in ms, length measurements (Stride Length; Stance Width) are in cm, and Stride Frequency is strides/s. Results from 38 mice (13 WT, 16 Het, and 9 I212F) are expressed as Mean ± SEM. See Table 1-Source Data File. ^1^WT, *Drd2*^+/+^; Het, *Drd2^+/I212F^;* I212F, *Drd2^I212F/I212F^* *p < 0.05; **p < 0.01; ***p < 0.001 compared to *Drd2*^+/+^ mice ^†^p < 0.05, compared to *Drd2^I212F/I212F^*

|  | <u>Swing Time</u> | <u>Propel Time</u> | <u>Brake Time</u> | <u>Propel/Brake</u> | <u>Stride Freq</u> | <u>Stride Length</u> | <u>Stance Width</u> |
| --- | --- | --- | --- | --- | --- | --- | --- |
| **Forelimbs** |  |  |  |  |  |  |  |
| **WT<sup>1</sup>** | 94 ± 4 | 93 ± 5 | 65 ± 4 | 1.63 ± 0.16 | 4.07 ± 0.09 | 6.07 ± 0.13 | 1.5 ± 0.07 |
| **Het** | 109 ± 2*** | 100 ± 3<sup>†</sup> | 62 ± 3 | 1.71 ± 0.09 | 3.78 ± 0.07** | 6.49 ± 0.15* | 1.6 ± 0.04 |
| **I212F** | 109 ± 3** | 114 ± 4** | 56 ± 3 | 2.19 ± 0.21* | 3.63 ± 0.06*** | 6.72 ± 0.10*** | 1.5 ± 0.06 |
| ANOVA | F(2,73)=9.9<br>p=0.0002 | F(2,70)=8.8<br>p=0.0004 | F(2,73)=1.7<br>p=0.19 | F(2,73)=3.5<br>p=0.035 | F(2,73)=7.8<br>p=0.0009 | F(2,73)=7.5<br>p=0.001 | F(2,35)=0.39<br>p=0.68 |
| **Hindlimbs** |  |  |  |  |  |  |  |
| **WT** | 98 ± 3 | 110 ± 5 | 36 ± 2 | 3.10 ± 0.14 | 4.20 ± 0.09 | 5.85 ± 0.11 | 3.0 ± 0.07 |
| **Het** | 107 ± 3 | 126 ± 3** | 40 ± 2 | 3.49 ± 0.23 | 3.76 ± 0.06*** | 6.54 ± 0.09*** | 2.8 ± 0.04 |
| **I212F** | 101 ± 3 | 136 ± 4*** | 41 ± 2 | 3.62 ± 0.32 | 3.71 ± 0.09*** | 6.67 ± 0.13*** | 2.9 ± 0.05 |
| ANOVA | F(2,73)=2.53<br>p=0.087 | F(2,73)=10.9<br>p<0.0001 | F(2,73)=1.19<br>p=0.31 | F(2,73)=1.26<br>p=0.29 | F(2,73)=12.2<br>p<0.0001 | F(2,73)=15.7<br>p<0.0001 | F(2,35)=1.97<br>p=0.154 |

### Striatal D2 Receptor Density

D2-I^212^F is expressed at 35-40% of the density of D2-WT after transfection of a given amount of DNA in HEK293 cells (Rodriguez-Contreras *et al.*, 2021; van der Weijden *et al.*, 2021a). We carried out radioligand binding studies with striatal tissue from a subset of the mice used for gait analysis to determine if D2 receptor density was also reduced in *Drd2^I212F^* mice. Genotype significantly affected the density of neostriatal D2 receptors (Figure 2; F(2,26)=20.61; p<0.0001). The density of receptors was decreased by 30% in *Drd2^+/I212F^* mice, and by 58% in *Drd2^I212F/I212F^* mice (Figure 2).

**Figure 2.**
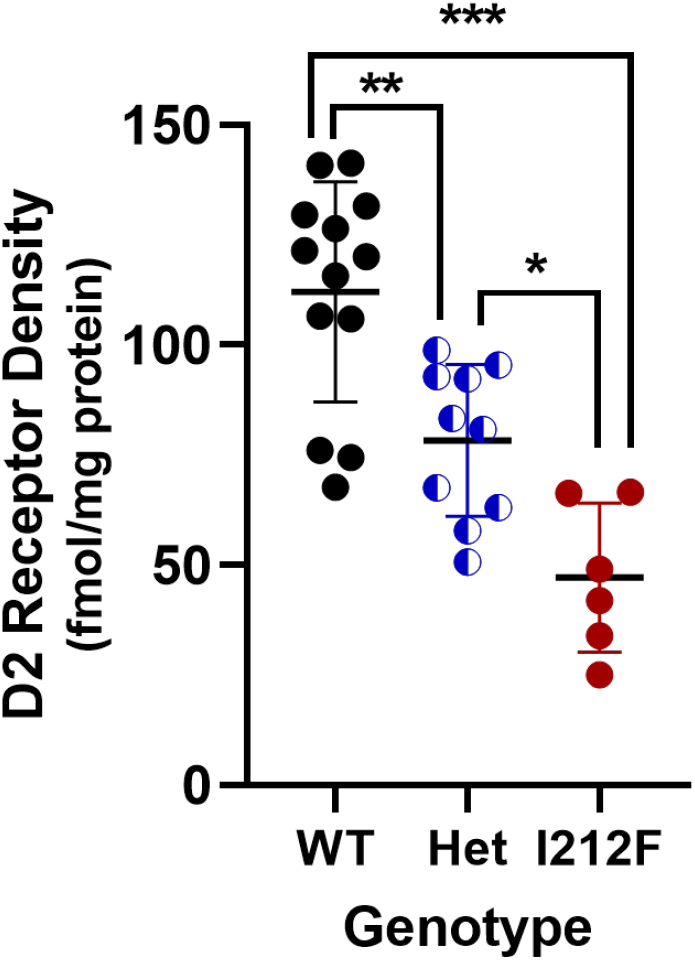
Striatal D2 receptor density. Results are shown for B_max_ values in striatal membranes from 10-12 month old *Drd2^+/+^* (WT), *Drd2^+/I212F^* (Het), and *Drd2^I212F/I212F^* (I212F) mice. Saturation binding analysis was carried out using the D2 antagonist radioligand [^3^H]spiperone, and B_max_ values were obtained by nonlinear regression. Mean values ± SD are shown. *p < 0.05; **p < 0.01; ***p < 0.001 for the indicated comparisons.

### Midbrain D2 Autoreceptors

We used *Drd2^I212F^* mice to confirm some of the results obtained after AAV-mediated expression of the variant in midbrain dopamine neurons (Rodriguez-Contreras *et al.*, 2021; van der Weijden *et al.*, 2021a). Midbrain slices prepared from mice of three genotypes (*Drd2*^+/+^ (WT), *Drd2^+/I212F^* (HET), and *Drd2^I212F/I212F^* (I212F)) were electrically stimulated (5 stimuli at 40 Hz) to elicit D2 receptor-GIRK IPSCs in response to somatodendritic dopamine release. Electrically evoked IPSCs from *Drd2^I212F^* mice were slow compared to those in preparations from wild type (Figure 3A). The amplitudes of evoked IPSCs were not significantly different among genotypes (Figure 3B: Kruskal-Wallis test, H = 2.93, p > 0.05). However, the duration of the IPSCs was increased in both *Drd2^I212F/I212F^* and *Drd2^+/I212F^* mice relative to controls (Figure 3C; one-way ANOVA, F = 192.7, p < 0.0001). We assessed the role of signal termination on this widening of the IPSC by calculating the tau of decay. *Drd2*^+/+^ and *Drd2^I212F/I212F^* mice were fit by a single exponential, with the latter showing a considerable slowing of the decay back to baseline (Figure 3D; Kruskal-Wallis test, H = 35.22, p < 0.0001). Interestingly, the evoked IPSCs in dopamine neurons from the *Drd2^+/I212F^* mice were fit by a double exponential (Figure 3D), one resembling *Drd2*^+/+^ cells (Het_Fast_) and one resembling *Drd2^I212F/I212F^* cells (Het_Slow_). Each component of this double exponential was similar to the tau measured in the corresponding homozygous genotype, reflecting the presence of distinct components arising from the two variants.

**Figure 3.**
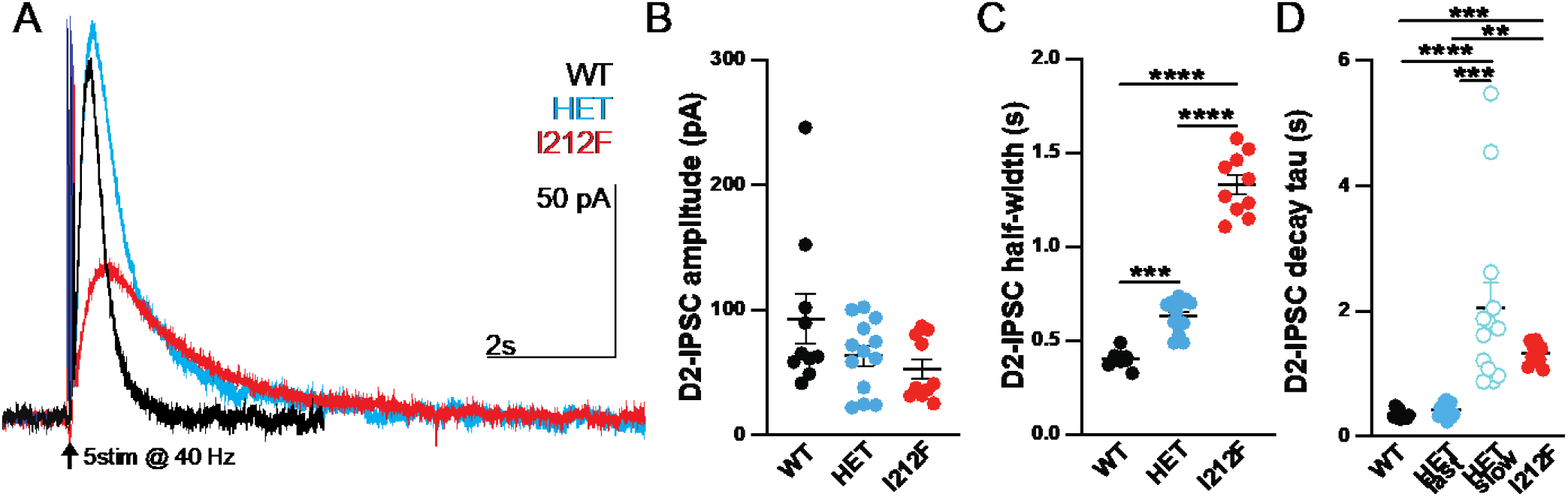
*Drd2^I212F^* mice exhibit a slowing of D2-autoreceptor IPSCs. **A**. Representative recordings of D2-IPSCs elicited by five electrical pulses delivered at 40 Hz in midbrain slides from *Drd2^+/+^* (WT, black), *Drd2^+/I212F^* (HET, cyan) and *Drd2^I212F/I212F^* (I212F, red) mice. **B**. Average amplitude of D2-IPSCs elicited with 5 pulses in each genotype. (WT = 93 ± 20 pA, HET = 63 ± 8 pA, and I212F = 53 ± 8 pA, Kruskal-Wallis test, H = 2.93, p > 0.05) **C**. Average width of the D2-IPSC measured at 50% of the peak (half-width) (WT = 0.40 ± 0.01 s, HET = 0.63 ± 0.03 s, and MUT = 1.33 ± 0.05 s. **D**. Tau values obtained with either a single (WT, I212F) or double (HET) exponential fit. *Drd2*^+*/I212F*^ mice displayed a two component decay, with a fast component resembling *Drd2*^+/+^ and a slow component resembling tau values for *Drd2*^I212F/I212F^ (WT = 0.3 ± 0.2 s; Het_Fast_, 0.43 ± 0.03 s; Het_Slow_, 2.1 ± 0.4 s; and I212F = 1.3 ± 0.1 s). Values plotted are the mean ± SEM of 10 cells/3 mice (WT), 13 cells/4 mice (HET), and 10 cells/3 mice (I212F), respectively.

We also examined the termination of signaling kinetics using exogenously applied dopamine (10 μM) and photolytic release of the competitive inverse agonist sulpiride from CyHQ-sulpiride (Asad *et al.*, 2020), since the amplitude and kinetics of D2-IPSCs may be influenced by changes in evoked dopamine release. Cells from all genotypes exhibited an outward current induced by 10 μM DA (Figure 4A) that returned to baseline following photolytic release of sulpiride (Figure 4B). The maximum current induced by 10 μM dopamine in cells from *Drd2^I212F/I212F^* animals was reduced to ~50% of that measured in cells from *Drd2*^+/+^ or *Drd2^+/I212F^* mice (Figure 4C, one-way ANOVA; F = 4.994, p < 0.05). Photolytic release of sulpiride produced a fast decay of signaling in cells from all three genotypes. However, as for the evoked IPSCs in Figure 3, the rate of decay was gene dosage dependent (Figure 4D, one-way ANOVA: F = 30.73, p < 0.0001).

**Figure 4.**
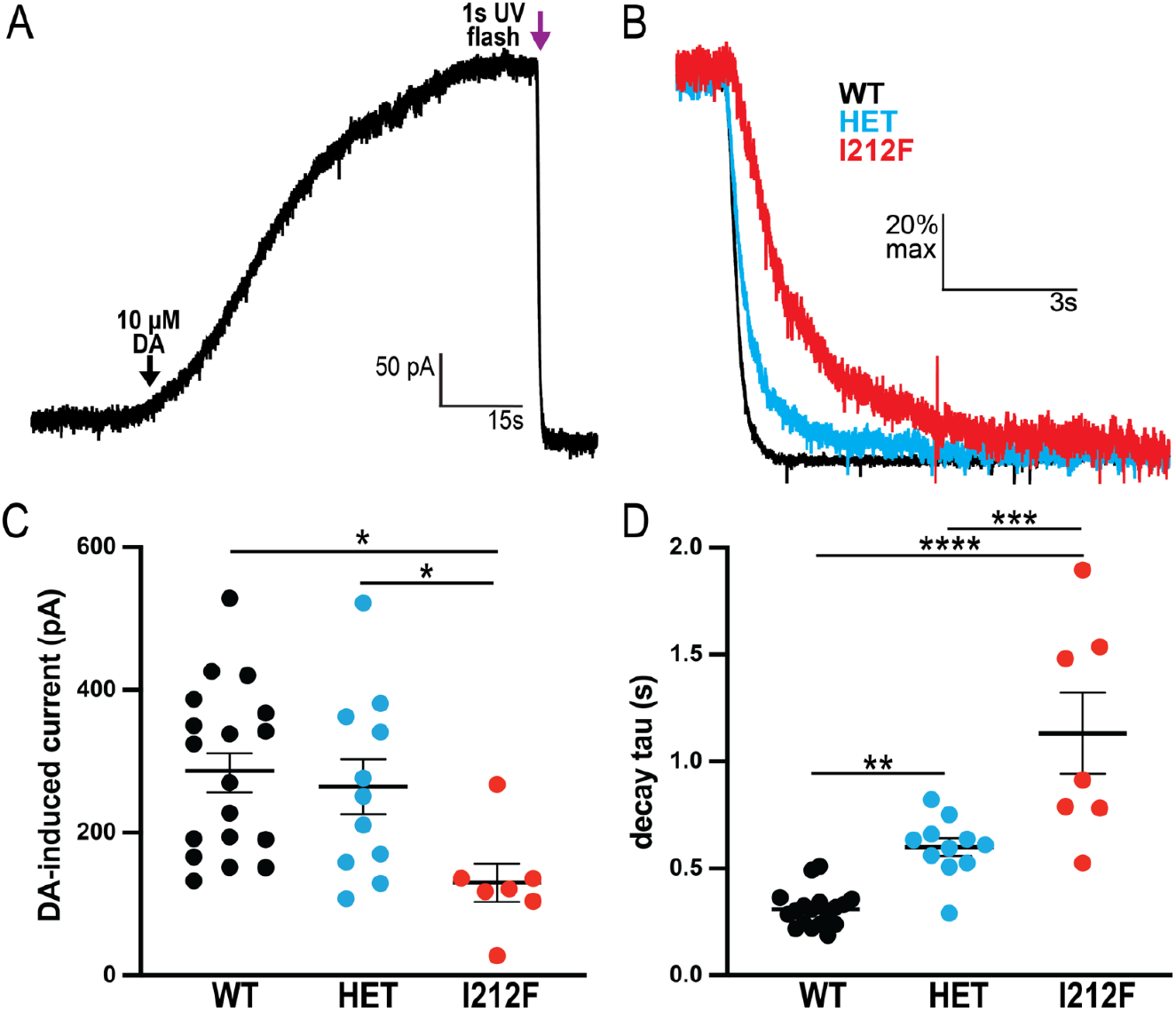
Prolonged decay kinetics in midbrain DA neurons from *Drd2*^I212F/I212F^ mice. **A**. Representative uncaging experiment in a WT mouse. CyHQ-sulpiride (15 μM) was recirculated for 5 min prior to the addition of 10 μM dopamine (black arrow). The resulting outward current was allowed to stabilize at a peak response before uncaging of sulpiride with a 1 s flash of UV light (purple arrow). **B**. Normalized (scaled-to-peak) representative D2 currents terminated by photolytic release of sulpiride in *Drd2^+/+^* (WT, black), *Drd2^+/I212F^* (Het, cyan), and *Drd2^I212F/I212F^* (I212F, red) animals. **C**. Amplitude of the current induced by 10 μM dopamine across genotypes. Homozygous mutants displayed a significant decrease compared to both WT and heterozygous animals (WT: 287 ± 27 pA, HET: 264 ± 38 pA, I212F: 130 ± 27 pA; Tukey’s test following one-way ANOVA; F = 4.994, p < 0.05; WT vs I212F q= 4.405, p = 0.0103; HET vs I212F q = 3.487, p= 0.0487). **D**. Quantification of the decay tau following photolytic release of sulpiride. Homozygous mutants exhibit the slowest kinetics, with heterozygous animals showing an intermediate slowing effect compared to WT animals (WT: 310 ± 20 ms, HET: 600 ± 42 ms, I212F: 1132 ± 191 ms; one-way ANOVA: F = 30.73, p < 0.0001; WT vs HET q= 4.536, p = 0.0081; WT vs I212F q= 11.04, p < 0.0001; HET vs I212F q = 6.583, p= 0.0001). Values plotted are mean ± SEM of 18 cells/6 mice (WT), 11 cells/4 mice (HET) and 7 cells/4 mice (I212F), respectively. * = p < 0.05, ** = p < 0.01, *** = p < 0.001, by Tukey’s post hoc test following one-way ANOVA.

### Postsynaptic D2 Receptors in Basal Forebrain

D2-I^212^F receptors expressed in HEK293 cells or dopamine neurons in the mouse midbrain differ in sensitivity to agonist and kinetics of GIRK regulation compared to D2-WT receptors (Figures 3 and 4; also Rodriguez-Contreras *et al.*, 2021; van der Weijden *et al.*, 2021a). Here, we used *Drd2^I212F^* mice to examine whether the kinetics and sensitivity of D2-I^212^F receptors also differ from D2-WT in D2-MSNs. An adeno-associated virus (AAV) encoding a GIRK2 channel and a tdTomato fluorophore was injected into both the dorsal striatum (DSt) and the nucleus accumbens shell (NAc) of *Drd2^I212F/I212F^* mice or their wild-type littermates (Figure 5A), as described previously (Marcott *et al.*, 2018; Gong *et al.*, 2021). Three weeks later, coronal brain slices containing the DSt and NAc were cut for electrophysiological recordings. Recordings were made in the presence of NMDA, GABAA, GABAB, muscarinic, and D1 receptor antagonists to isolate D2 receptor mediated GIRK2 currents. In whole-cell voltage clamp, a single electrical stimulus evoked D2 receptor-mediated IPSCs (D2-IPSCs) in D2-MSNs (Figure 5B). We observed that the amplitude of D2-IPSCs was smaller in both the DSt and NAc of *Drd2^I212F/I212F^* mice compared to littermate controls (Figure 5C). Similar to somatodendritic IPSCs recorded from dopamine neurons, electrically evoked D2-IPSCs in both DSt and NAc D2-MSNs from *Drd2^I212F^* mice were slower to activate (Figure 5D) and were slower to decay than littermate controls (Figure 5E). The percent changes between D2-IPSCs recorded in *Drd2*^I212F^ vs. D2-WT mice were similar in the DSt and NAc (10 −90% rise: DSt = 69% increase vs. D2-WT; NAc = 61% increase vs. D2-WT; tau decay: DSt = 650% increase vs. D2-WT; NAc = 573% increase vs. D2-WT), indicating that our previous observation that D2 receptor kinetics are slower in the NAc than in the DSt (Marcott *et al.*, 2018) also holds for mice expressing D2-I^212^F. The results suggest that the D2-I^212^F mutation causes slower D2 receptor signaling kinetics in both the DSt and the NAc.

**Figure 5.**
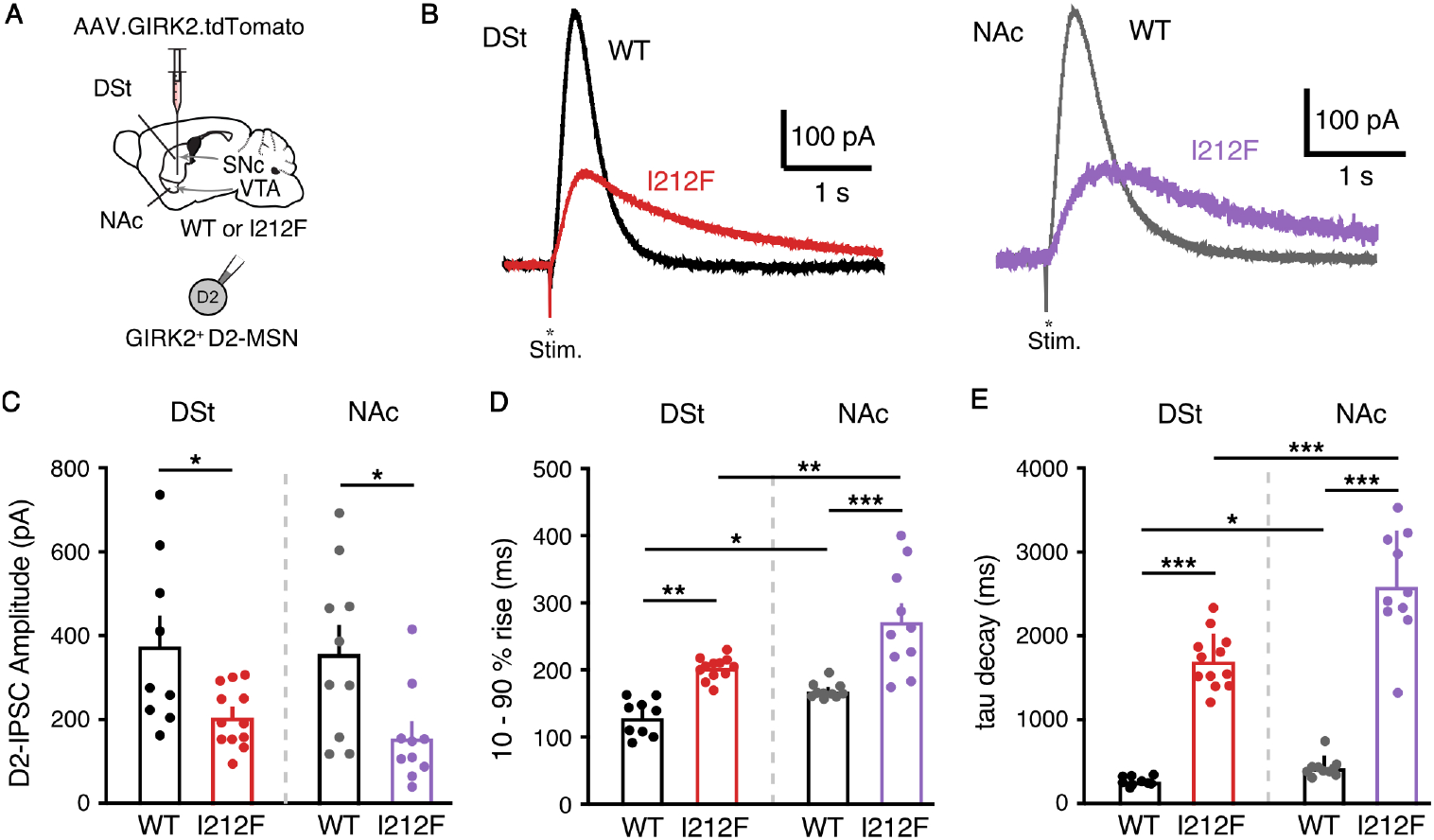
Characterization of D2-IPSCs in striatal medium spiny neurons. **A**. Injection schematic of AAV9.hSyn.tdTomato.GIRK2 into the NAc (medial shell) and DSt (dorsomedial) of the *Drd2^+/+^* (WT) or *Drd2^I212F/I212F^* (I212F) mice. **B**. Representative traces of electrical stimulation-evoked D2-IPSCs from the DSt and NAc of WT and I212F mice. **C**. Quantification of the D2-IPSC amplitudes (DSt: D2-WT = 377 ± 67 pA, n = 9; D2-I^212^F = 206 ± 21 pA; n = 12; NAc: D2-WT = 358 ± 64 pA, n = 10; D2-I^212^F = 156 ± 36 pA, n =10). **D**. Quantification of 10 −90% rise time of D2-IPSCs (DSt: D2-WT = 120 ± 6 ms; D2-I^212^F = 203 ± 5 ms; NAc: D2-WT = 169 ± 4 ms; D2-I^212^F = 272 ± 25 ms; one-way ANOVA: F = 87.21, p < 0.001. **E**. Quantification of tau of decay of D2-IPSCs (DSt: D2-WT = 261 ± 14 ms; D2-I^212^F = 1702 ± 95 ms; NAc: D2-WT = 453 ± 34 ms; D2-I^212^F = 2595 ± 203 ms; one-way ANOVA: F = 23.78, p < 0.001. **C-E**: Summary data are mean ± SEM from 9 cells (DSt) and 10 cells (NAc)/6 mice (D2-WT), or 12 cells (DSt) and 10 cells (NAc)/7 mice (D2-I^212^F). * = p < 0.05, ** = p < 0.01, *** = p < 0.001 by Mann-Whitney U test (**C**) or Sidak’s multiple comparisons test (**D-E**).

Given that D2-IPSCs evoked by the release of dopamine differed in *Drd2^I212F/I212F^* mice, we next examined if the sensitivity of D2 receptor signaling also differed. Concentration-response relationships for dopamine in the DSt and NAc were constructed by measuring D2 receptor mediated GIRK2 currents evoked by bath application of dopamine in the presence of cocaine (10 μM) to block dopamine reuptake (Figure 6A). Similar to our previous findings (Marcott *et al.*, 2018; Gong *et al.*, 2021), the concentration of dopamine needed to achieve 50% of the maximal effect (EC_50_) in D2-WT mice was significantly lower in the NAc than the DSt (Figure 6B and C), confirming that D2 receptors in the NAc have a higher sensitivity for dopamine than in the DSt. The D2-I^212^F mutation led to a selective increase in the sensitivity of D2 receptors in the DSt, as reflected in a decrease in the EC_50_ value, but had no effect on the sensitivity of D2R signaling in the NAc (Figure 6C). The mutation also decreased the amplitude of the outward current evoked by a saturating concentration of dopamine (300 μM) (Emax) in both regions, an effect that was greater in the DSt than the NAc (Figure 6D). As the difference in D2 receptor sensitivity between the DSt and NAc in wild type mice results from differential coupling to Gα_o_ over Gα_i_ in the NAc (Marcott et al., 2018; Gong et al., 2021), the present results are consistent with our prior work demonstrating that agonist potency at D2-I^212^F is similar for activation of Gα_o_ and Gα_i_ (Rodriguez-Contreras *et al.*, 2021).

**Figure 6.**
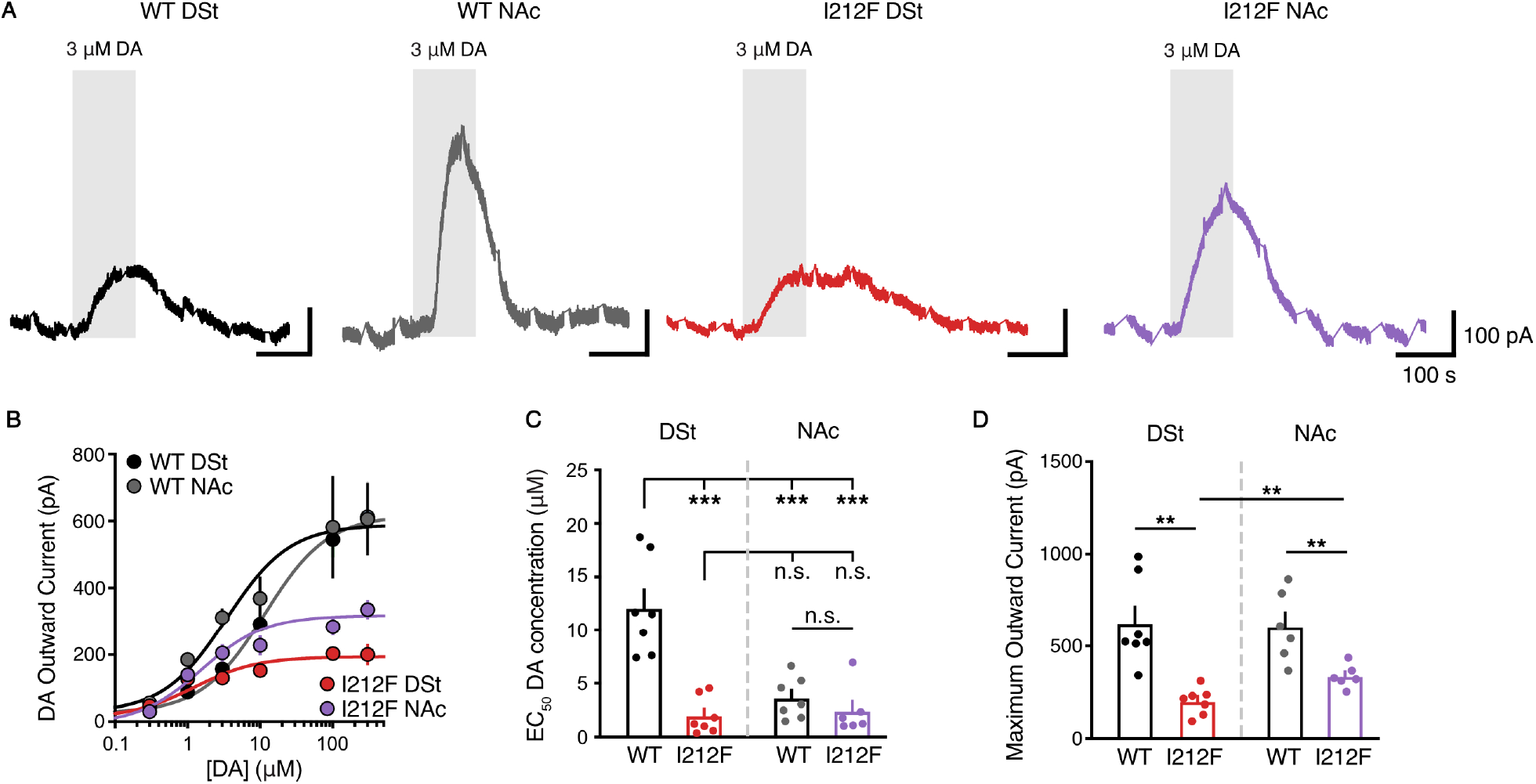
Dopamine potency in striatal D2-MSNs. **A**. Representative traces after bath application of dopamine (DA, 3 μM) depict dopamine-induced D2R-mediated outward currents from D2-MSNs in the DSt and NAc from *Drd2^+/+^* (WT) or *Drd2*^I212F/I212F^ (I212F) mice. Recordings were performed in the presence of cocaine (10 μM) to block dopamine uptake. D2-IPSCs were evoked once per minute and have been blanked for clarity. **B**. Dopamine concentration-response relationships for D2 receptor-mediated outward GIRK2 current. **C**. EC_50_ values calculated from dopamine concentration-response curves (shown in **B**) (DSt: D2-WT = 12.1 ± 1.7 μM; D2-I^212^F = 2 ± 0.7 μM; NAc: D2-WT = 3.6 ± 0.7 μM; D2-I^212^F = 2.4 ± 0.9 μM; Tukey’s test following one-way ANOVA: F = 18.87, p < 0.001). Summary data are mean ± SEM from 7 repetitions for D2-WT (DSt and NAc) and D2-I212F (DSt) and 6 repetitions for D2-I212F (NAc). **D**. Maximum outward currents evoked by 300 μM dopamine (from **B**) (DSt: D2-WT = 624 ± 89 pA, D2-I^212^F = 201 ± 27 pA; NAc: D2-WT= 606 ± 78 pA, D2-I^212^F = 336 ± 25 pA). Summary data are mean ± SEM of 6-7 cells from each brain region in 6-7 mice of each genotype, with statistical comparisons by Mann-Whitney test n.s. = p > 0.05, ** = p < 0.01, *** = p < 0.001.

We reported previously that chronic cocaine exposure selectively decreases D2 receptor sensitivity in the NAc via the reduction of Gα_o_ levels to gate cocaine-conditioned behaviors, whereas D2 receptor sensitivity for the Gα_i_-mediated response in the DSt is unaffected by prior cocaine exposure (Gong *et al.*, 2021).

Because agonist potency at D2-I^212^F is similar for Gα_o_ and Gα_i_, we predicted that prior cocaine exposure would have no effect on the potency of dopamine in DSt or NAc of *Drd2^I212F/I212F^* mice. To test this, *Drd2^I212F/I212F^* mice were treated with cocaine for 7 days (20 mg/kg intraperitoneally [i.p.]) (Figure 7A). As predicted, this cocaine exposure had no effect on D2 receptor sensitivity to dopamine in either the DSt or NAc of *Drd2^I212F/I212F^* mice (Figure 7B and C). Interestingly, chronic exposure to cocaine selectively reduced the outward current evoked by a saturating concentration of dopamine (300 μM) in the NAc of *Drd2^I212F/I212F^* mice, without having an effect in the DSt (Figure 7D). In wild type mice, prior cocaine treatment did not change the maximum response in either DSt or NAc (Gong *et al.*, 2021).

**Figure 7.**
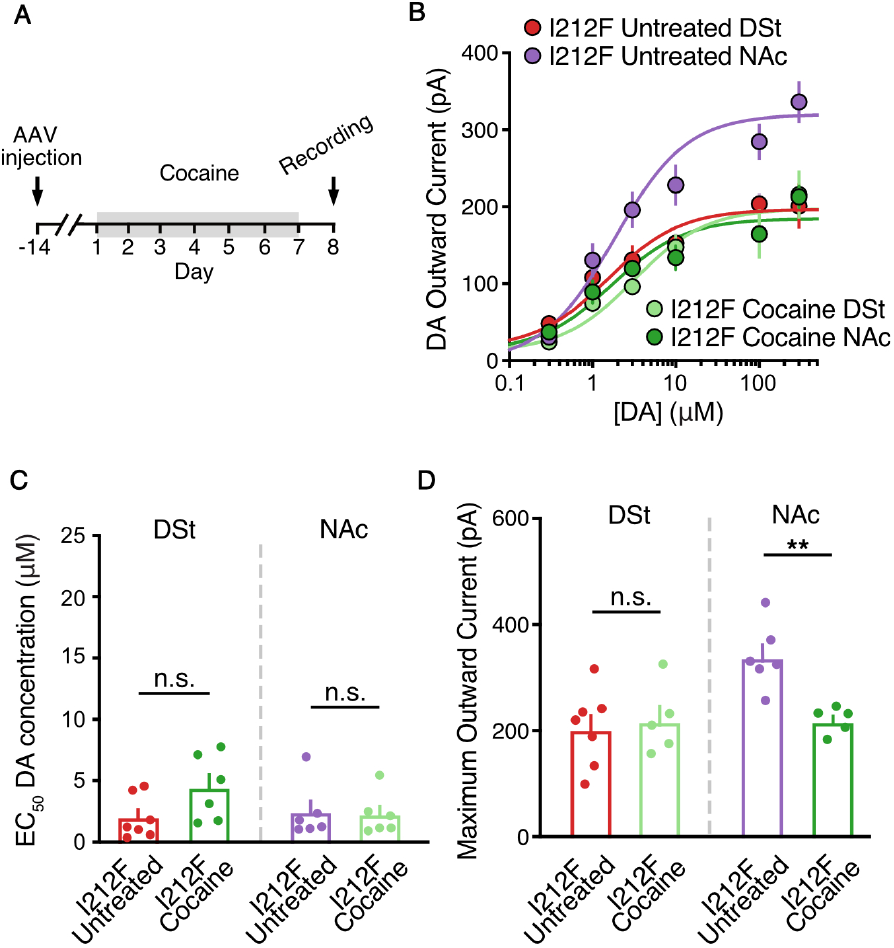
Effect of prior cocaine treatment. **A**. Schematic of the timeline of AAV.GIRK2 injection and 7-day cocaine administration. **B**. Dopamine concentration-response relationships for D2 receptor-mediated outward GIRK2 current from D2-MSNs in the DSt and NAc from *Drd2^I212F/I212F^* (I212F) mice after 7 days cocaine exposure (Cocaine) or from control animals (Untreated; from Figure 6B). **C**. EC_50_ values from (B). **D**. Maximum outward currents evoked by 300 μM dopamine from (B). Summary data are mean ± SEM. n.s. = p > 0.05, ** = p < 0.01; Mann-Whitney

## DISCUSSION

Human carriers of the dopamine D2 receptor mutation c.634A>T, p.Ile212Phe have a hyperkinetic movement disorder characterized by symptoms of chorea and/or dystonia that appear in adolescence and worsen with increasing age (van der Weijden *et al.*, 2021a; van der Weijden *et al.*, 2021b). This novel mutation in exon 5 of *DRD2* changes residue 212 at the cytoplasmic face of transmembrane domain 5 of the D2 receptor from Ile to Phe (Figure 1A).

The clinical phenotype associated with heterozygosity for D2-I^212^F has features in common with other genetic disorders characterized by childhood-onset chorea, frequently accompanied by dystonia and/or myclonus. One such disorder, called either Familial Dyskinesia with Facial Myokymia (FDFM; Fernandez *et al.*, 2001; Chen *et al.*, 2012) or ADCY5-related dyskinesia (MIM: 606703; Carecchio *et al.*, 2017), is caused by activating mutations of adenylyl cyclase 5 (Chen *et al.*, 2014; Doyle *et al.*, 2019), a striatum-enriched form of adenylyl cyclase that mediates D1 dopamine receptor stimulation and D2 receptor inhibition of adenylyl cyclase in medium spiny neurons of the neostriatum and nucleus accumbens (Lee *et al.*, 2002). It is noteworthy that a second pathogenic *DRD2* mutation, which changes a residue at the cytoplasmic face of transmembrane domain 6, also causes a hyperkinetic movement disorder with an early childhood onset, and that one of the carriers of this mutation was part of a sample population that had previously been tested for *ADCY5* mutations (Mencacci *et al.*, 2021; van der Weijden *et al.*, 2021b).

Similarly, many mutations of *GNAO1* (Gα_oA_) cause a syndrome called “neurodevelopmental disorder with involuntary movements” (MIM: 617493; Ananth *et al.*, 2016). Loss-of-function mutations of Gα_oA_ are associated with epilepsy, whereas activating mutations are associated with early-onset dyskinesia including choreatic features (Feng *et al.*, 2018). Because Gα_o_ mediates many effects of D2 receptor signaling (Jiang *et al.*, 2001; Marcott *et al.*, 2018), it might be expected that activating mutations of the two proteins would cause overlapping phenotypes.

Our prior work using heterologous expression in HEK 293 cells determined that basal G protein-mediated signaling and agonist potency are enhanced for D2-I^212^F, with the magnitude of effects depending on the Gα subtype, whereas agonist-induced binding of arrestin to D2-I^212^F is decreased. Molecular dynamics simulations (Rodriguez-Contreras *et al.*, 2021) suggest that these functional changes may be the result of weakening an intramolecular salt bridge that maintains some G protein-coupled receptors in an inactive conformation (Ballesteros *et al.*, 2001). We confirmed the effects of the mutation on G protein-mediated signaling in midbrain dopamine neurons after viral expression of D2-I^212^F, and also determined that a consistent difference between the wild type D2 receptor and D2-I^212^F is that the rate at which Gβγ-mediated activation of GIRKs is terminated is many-fold slower for the pathogenic variant (Rodriguez-Contreras *et al.*, 2021; van der Weijden *et al.*, 2021a).

We have now produced a mouse constitutively expressing D2-I^212^F to confirm the pathogenicity of this variant and as a tool to further evaluate the functional consequences of the mutation. Using the DigiGait system, we found that a number of gait characteristics were altered in *Drd2^I212F^* mice tested at 10-12 months of age. *Drd2^212F/I212F^* mice took longer to swing forelimbs forward, took longer strides with hind- and forelimbs, and had decreased stride frequency for both hind- and forelimbs. The gait analysis also estimates the proportion of the time that paws are in contact with the treadmill that is spent propelling the mouse forward vs. braking. Homozygous mutant mice spent more time in hindlimb and forelimb propulsion, and had an increased ratio of forelimb time spent in propulsion to time spent braking. Importantly, considering that all humans known to have this variant are heterozygous at the *DRD2* locus, *Drd2^+/I212F^* mice exhibited most of the same gait abnormalities, supporting the pathogenic role of the D2-I^212^F variant in humans.

In HEK 293 cells, D2-I^212^F is expressed at a lower density than D2-WT after transfection of identical amounts of DNA (van der Weijden *et al.*, 2021a). Similarly, in *Drd2^I212F^* mice, the density of neostriatal D2 receptors was decreased in both heterozygous and homozygous mutant mice, with the magnitude of the decrease dependent on the number of *Drd2^I212F^* alleles. This may be a consequence of receptor constitutive activity, as constitutively active G protein-coupled receptors have often been shown to be less stable and therefore degraded more rapidly (Gether *et al.*, 1997; Rasmussen *et al.*, 1999; Alewijnse *et al.*, 2000), although it is also possible that maturation and trafficking of the receptor to the plasma membrane is delayed by the mutation. It was previously reported that striatal binding potentials for the D2/D3 receptor positron emission tomography ligand [^11^C]raclopride in three subjects heterozygous for *DRD2*^I212F^ were within the normal range (van der Weijden *et al.*, 2021a). The modest decrease that the results from *Drd2^+/I212F^* mice suggest should be found in heterozygous subjects may be difficult to detect given the low number and the age range of the subjects, and in the context of normal aging-related declines in D2 receptor density.

We confirmed the slow kinetics of D2-I^212^F-mediated GIRK conductances seen in viral overexpression experiments using midbrain slices from homozygous *Drd2^I212F^* mice. Importantly, the two components mediated by D2-WT and D2-I^212^F in the heterozygous mice combined to produce GIRK currents that were substantially prolonged compared to those in wild type mice, whether assessed by the decay of evoked IPSCs or by photolytic release of sulpiride.

Slow kinetics are not limited to D2-I^212^F receptors expressed in dopamine neurons, as we observed similar results in striatal D2 receptor-expressing MSNs. The finding that sulpiride-induced termination of the GIRK response to bath-applied dopamine or in the absence of dopamine is slower for D2-I^212^F-mediated currents (present results and Rodriguez-Contreras *et al.*, 2021) indicates that the slow kinetics probably reflect post-synaptic mechanisms. Prolonged GIRK activation suggests that activation of the G protein heterotrimer by D2-I^212^F results in availability of free of Gβγ that is prolonged relative to activation that is mediated by D2-WT, but the mechanistic basis for prolonged elevation of Gβγ is yet to be determined. Although one hypothesis could be that D2-I^212^F desensitizes more slowly because of decreased arrestin binding, we found no evidence of reduced desensitization in response to bath-applied quinpirole (van der Weijden *et al.*, 2021a). A second hypothesis is that D2-I^212^F receptors exhibit higher affinity for activating ligands, slowing the competitive inhibition by sulpiride, although our previous finding that decay is slower for D2-I^212^F even in reserpine-treated slices argues against an exclusive role for slow unbinding of dopamine (Rodriguez-Contreras *et al.*, 2021). Experiments using sulpiride uncaging with agonists of varying affinities will be required to definitively distinguish slow unbinding from downstream signaling changes (Condon *et al.*, 2021).

Prior work has shown that differential coupling to G protein subtypes leads to higher potency D2 receptor signaling in D2-MSNs of the nucleus accumbens than the dorsal neostriatum, and that the high potency signaling is lost after repeated daily treatment with cocaine (Marcott *et al.*, 2018; Gong *et al.*, 2021). The high- and low-potency signaling in untreated mice reflects D2 receptor coupling to Gα_o_ and Gα_i_, respectively, and the loss of accumbal high-potency signaling results from a cocaine-induced reduction in the abundance of Gα_o_. In the current study, we demonstrated that the potency of dopamine is similar in NAc and DSt and unaffected by prior cocaine treatment in homozygous *Drd2*^I212F^ mice. In these mice, however, cocaine treatment selectively decreased the maximal response to dopamine in NAc, a response that was not previously observed in wild type mice (Gong *et al.*, 2021). It may be that the combined effect of the cocaine-induced decrease in Gα_o_ and the lower expression of D2-I^212^F is to decrease the maximal response, instead of decreasing potency as observed for D2-WT (Gong *et al.*, 2021). Determining the consequences of these differing responses to cocaine treatment for behavior such as cocaine-conditioned place preference requires further study. We anticipate that this novel mouse model will be useful for such studies as well as for investigations into the mechanisms and treatment of early onset hyperkinetic movement disorders.

## MATERIALS AND METHODS

### Animals

All studies were conducted in accordance with the Guide for the Care and Use of Laboratory Animals established by the National Institutes of Health. Protocols were approved by Institutional Animal Care and Use Committees at the VA Portland Health Care System, Oregon Health & Science University (OHSU), and the University of Colorado School of Medicine. All mice were allowed ad-lib access to food and water and were maintained on a 12h light/dark cycle in a climate-controlled facility.

### Design and Generation of *Drd2^+/I212F^* mice

Knock-in *Drd2^+/I212F^* mice were produced via the electroporation of one-cell-stage C57BL6/NJ mouse embryos using a NEPA 21 electroporator (NEPA GENE Co. Ltd., Chiba, Japan) as described (Teixeira *et al.*, 2018). Ribonucleoprotein complexes of SpCas9 protein (NEB; Ipswitch, MA), ssODN template and gRNA were prepared with final concentration 200 μg/μl, 100 μg/μl and 200 μg/μl respectively. The ssODN template TCCATCGTCTCGTTCTACGTGCCCTTCATCGTCACaCTGCTGGTCTATATCAAA***T***TCTA CATC<u>GT*A*C</u>TaCGCAAGCGTCGGAAGCGGGTCAACACCAAGCGTAGCAGCCGAGCT incorporates the desired ***T*** change (bold, italic) in exon 5 of the *Drd2* gene to introduce the I^212^F mutation, a synonymous *A* mutation (italic in the underlined sequence) to introduce an RsaI site for genotyping purpose, and two additional synonymous mutations (lowercase) to remove Cas9 PAM sequences. This DNA template was synthesized by Integrated DNA Technologies (Coralville, IA). The gRNA (UGACCCGCUUCCGACGCUUG) was synthesized as a chemically modified CRISPR gRNA by Synthego Corporation (Redwood City, CA). After electroporation, embryos were transplanted into pseudopregnant recipient CD-1 female mice, and founders from these litters were identified as described below.

Putative F0 founders were genotyped by PCR/digestion and Sanger sequencing (see details below). Selected F0 founders were also analyzed by Sanger sequencing for potential modification of CRISPOR-predicted off-target (OT) sites (see below). Founders were crossed with C57BL/6JN mice, and F1 pups were genotyped by PCR/digestion and DNA sequencing.

### Analysis of target site and offspring genotyping

Mice potentially containing the knock-in mutation were initially screened for DNA sequence changes in *Drd2* exon 5 by PCR and RsaI digestion analysis as described below. The I^212^F mutation introduces an ApoI restriction site that was also used for genotyping with ApoI-HF. Sequence-specific forward and reverse primers (Supplementary File 1 and Figure 1C) for amplifying exon 5 and the introns/exon 5 junctions, were designed against the reference *Mus musculus* genome, strain C57BL/6NJ (GeneBank assembly accession: GCA_001632555.1). Extraction of genomic DNA was performed using the HotSHOT method (Truett *et al.*, 2000). Briefly, tail snip samples were heated in 100 ul of alkaline buffer (25 mM NaOH, 0.2 mM EDTA) at 95°C for 25 minutes, and neutralized with 100 ul of 40 mM Tris.HCl, pH 5. Two-three μl of supernatant were used per 20 ul PCR reaction. PCR was performed using the HotStarTaq Master Mix Kit (Qiagen; Germantown Road, MD) under the following conditions: 95°C for 3 min, 35 cycles at 95°C for 15s, 58°C for 15s, 72°C for 30s and a final extension step at 72°C for 5 min. PCR products were digested with RsaI or ApoI-HF (NEB Inc; Ipswich, MA) and were separated on 1.5% agarose gels. Mice harboring the on-target *Drd2*-exon 5 knock-in allele were identified based on fragment size after digestion of the PCR product. For example, PCR amplification using *Exon5-F1* and *Exon5-RC1* primers (Supplementary File 1) generated a 376-bp product (Figure 1-figure supplement 1A). After restriction digestion, the mutated allele generated two fragments of ~0.19 kb differing from the undigested wild type (WT) fragment of ~0.38 kb (Figure 1-figure supplement 1B). F1 offspring from selected Founders crossed with C57BL/6J (WT) mice were genotyped following the same procedure.

### Mutation detection by Sanger sequencing

To confirm positive F0 heterologous-genotyped mice and their F1 offspring, the 181-bp and 376-bp amplicon products (Figure 1-figure supplement 1A) were purified using the Monarch PCR & DNA Cleanup Kit (New England Biolabs, Ipswich, MA) and sent for Sanger DNA sequencing at the OHSU Vollum Sequence Core (Portland, OR). To check the integrity of both WT and mutated DNA allelic strands, the 376-bp amplicon containing *Drd2*-exon5 and introns/exon 5 junctions from several F0 and F1 mice were cloned into TOPO-TA Cloning vector (Invitrogen by Thermo Fisher Scientific; Waltham, MA) according to manufacturer’s instructions. Plasmids were purified using the Wizard Plus mini-prep DNA purification system (Promega, Madison, WI) and screened for WT- and I^212^F-mutated allele strands by dual digestion analysis using EcoRI digestion (New England Biolabs, Ipswich, MA) to release the insert, and RsaI or ApoI-HF digestion for selecting WT versus mutated strand clones. Positive clones for each allelic strand per mouse were sent for DNA sequencing using the universal T7-Forward oligo.

### Molecular characterization of *Drd2* exon 5 in *Drd2^I212F^* mice

In lineage **A**, digestion of the PCR amplicon with ApoI-HF supported the heterozygosity of the I^212^F mutation. Direct DNA sequencing of the amplicon confirmed the presence of the I^212^F mutation and showed no mixed sequence reads in the F0-429 mouse (Figure 8A, Top). Cloning and sequencing of both alleles confirmed the heterozygosity of the I^212^F mutation in this founder. However, the F0-429 mouse was homozygous for the RsaI site (Figure 8A, Bottom), in agreement with the results obtained by restriction enzyme analysis and direct DNA sequencing of the PCR amplicon (Figure 8A, Top). No other modifications were observed in this founder. The positive F1 offspring of this lineage (five females and two males) were heterozygous for the I^212^F mutation and RsaI site by both restriction enzyme analysis and DNA sequencing (example in Figure 8B).

**Figure 8.**
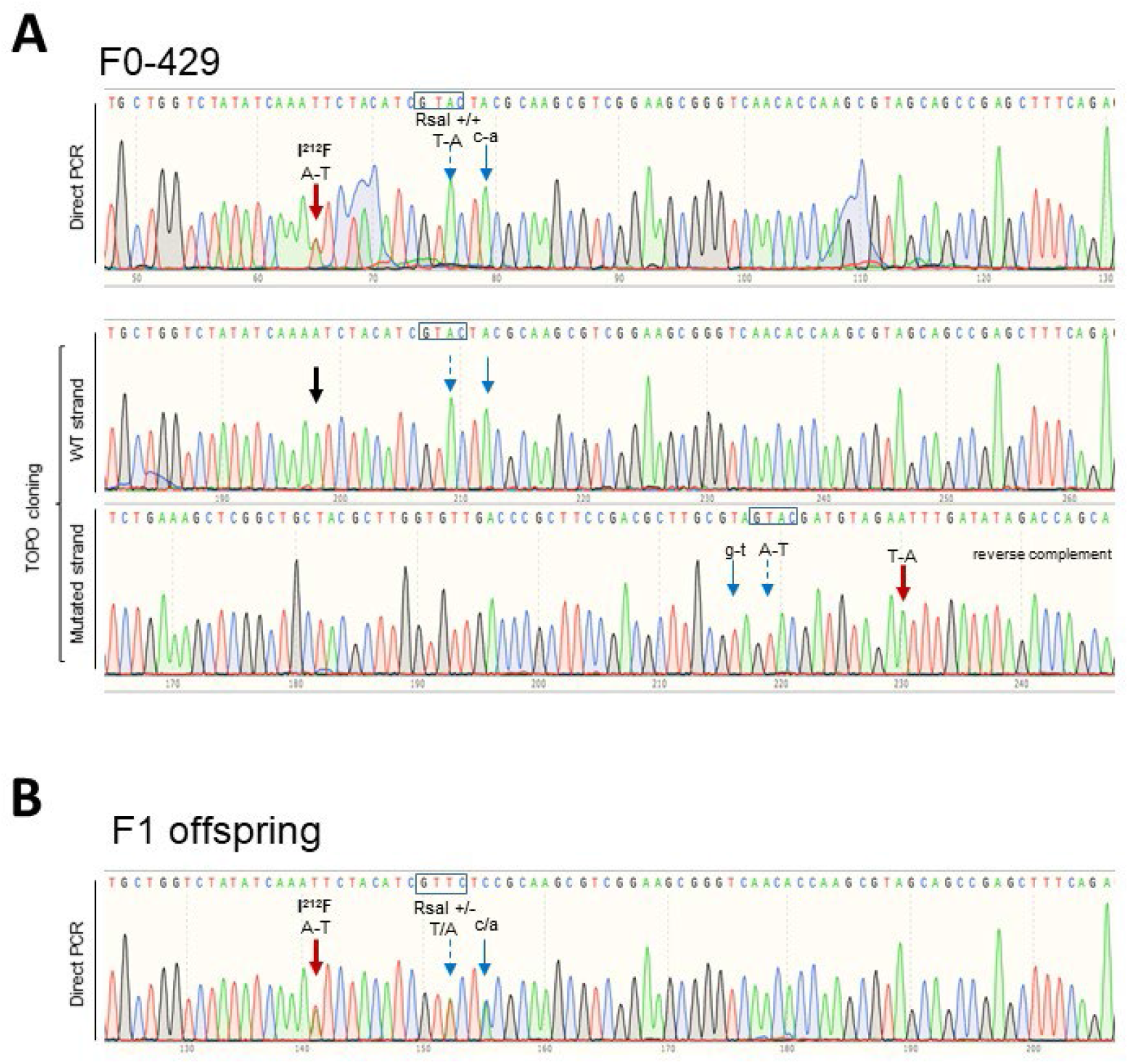
Production of *Drd2^I212F^* mice. **A**, Chromatograms showing the Sanger sequencing confirmation of the c.634A>T; p.Ile212Phe mutation in Founder F0-429 (lineage **A**). <u>Top</u>, 181-bp product obtained by PCR amplification of gDNA sample using the Exon5-F2 and Exon5-RC2 primer pair was purified and sent for sequencing. <u>Bottom</u>, 376-bp product amplified by PCR using the Exon5-F1 and Exon5-RC1 primer pair was cloned into TOPO-TA vector (Invitrogen), and plasmids were sent for sequencing. For the mutated strand, the reverse-complement sequence is shown. The Black arrow indicates the location c634A in the wild type strand. However, F0-429 was homozygous for both nucleotide changes introduced downstream of the c.634A>T position. **B**, Representative chromatogram showing Sanger sequencing of the 181-bp amplicon from an F1 mouse of lineage **A**. Founder F0-429 (male) was crossed with a C57BL/6NJ female (WT) mouse. F1 offspring (nine females and five males) were genotyped by PCR and restriction analysis. Sanger sequencing of 181-bp amplicons from positive F1 mice (five females and two males) confirmed that they were heterozygous for all designed nucleotide changes. Wide-red arrows in both panels point to the A-T change to generate the *Drd2*^I212F^ variant; blue arrows show synonymous nucleotide changes introduced downstream of the c.634A>T position. The nucleotide change in uppercase introduced to generate an RsaI site is indicated by a dashed-line arrow and its site (GTAC) is shown in a box, whereas the nucleotide change in lowercase was introduced to remove a gRNA PAM sequence. The molecular characterization of founder F0-421 (lineage **B**) and its F1 offspring is shown in Figure 8 - figure supplement 1.

In Lineage **B**, we found that founder F0-421 contains the expected I^212^F-mutation as well as the introduced RsaI site; however, mixed sequence reads were observed by direct DNA sequencing of the PCR product (Figure 8 – figure supplement 1A - Top). Analysis of the two alleles by TOPO-TA cloning and DNA sequencing revealed that one allele was WT for the targeted mutation site but had a 9-nt deletion downstream of that I^212^ site, whereas only the expected nucleotide changes, with no deletions/insertions, were present in the I^212^F-mutated allele (Figure 8 – figure supplement 1A - Bottom). We cloned and analyzed both *Drd2-exon* 5 alleles from its positive F1 offspring (two females and 5 males). As expected, WT-strands had no genetic modifications (Figure 8 – figure supplement 1B), whereas the mutated strands have only the nucleotide changes described in Figure 1C. Both *Drd2*^I212F^ lineages **A** and **B** (from founders F0-429 and F0-421, respectively) were maintained as heterozygous breeding colonies at the VA Portland Health Care System (VAPORHCS). In addition, mice of lineage **A** were maintained at the University of Colorado Anschutz Medical Campus.

### Analysis of off-target sites

The *Drd2*-exon 5 sequence (CM004223.1:49596634-49596825, strain +) was entered in the web-based tool CRISPOR (www.crispor.tefor.net; Concordet and Haeussler, 2018) to predict potential off-target (OT) sites for the gRNA. No off-target locus with less than four nucleotide mismatches to the selected gRNA (UGACCCGCUUCCGACGCUUG) was found in the C57BL/6J reference genome. To identify any OT mutations in the founders for the two lineages, PCR reactions were carried out using primer pairs designed to flank the potential OT1 to OT4 sites (Supplementary File 1). Three to four independently generated PCR products for each OT1-4/founder were purified using the Monarch PCR & DNA Cleanup Kit (NEB Inc.) and sent for Sanger sequencing at the OHSU Vollum Sequence Core.

The gRNA UGACCCGCUUCCGACGCUUG sequence had minimal predicted OT effects. Five sites with 4-nucleotide mismatches were identified (Supplementary File 2). However, only OT1 (CFD score 0.45918) and OT2 (CFD score 0.13333) sites had a Cutting Frequency Determination (CFD) score higher than 0.1 (Doench *et al.*, 2016). For OT3-5 sites, the CFD scores were 0.015278, 0.01500 and 0.00824, respectively (Supplementary File 2). Despite the low CFD score, we characterized OT4 because it is on the same chromosome as *Drd2* and therefore an unintended mutation would not be quickly eliminated in subsequent generations, and also characterized OT3 because its CFD score was slightly higher than OT4. To identify any OT mutagenesis/alteration effects in the F0-421 and F0-429 mice, PCR reactions were carried as described above. We found no modifications in the studied off-target sequences, when compared with the reference *Mus musculus* genome, strain C57BL/6NJ.

### Gait analysis

We used the DigiGait system (Mouse Specifics, Inc., Framingham, MA) to quantify gait abnormalities in *Drd2^I212F^* mice. DigiGait uses a variable-speed motorized transparent treadmill belt and a ventral video camera capturing 150 frames/s to calculate over 35 indices of gait for each limb. Each mouse was placed in a Plexiglas compartment that sits on top of the transparent treadmill and is illuminated from above and below. The mouse was allowed to explore the compartment for several minutes, then the camera was turned on and the treadmill was started at 24 cm/s. After 4-5 s the treadmill was turned off, the video clip was saved, and the mouse was returned to its home cage. Videos were processed by an individual blind to mouse genotype.

### D2 receptor radioligand binding

To measure membrane expression of striatal D2 receptors, pooled striata from each mouse were homogenized for 10 sec in 4 ml of Tris-buffered saline (TBS: 50 mM Tris, 120 mM NaCl, pH 7.4) using a Polytron homogenizer (Brinkmann Instruments, Westbury, NY), then centrifuged at 17,000 × *g* at 4°C for 20 min. The resulting pellet was resuspended in 4 ml TBS + 2 mM Na-EDTA, incubated for 30 min at 25°C, and centrifuged again at 17,000 × *g* for 20 min. The final pellet was resuspended in 3 ml TBS. Protein determination was performed using the BCA Protein Assay Kit (Thermo Scientific Inc; Waltham, MA). Saturation analyses were carried out by preparing ~0.4 nM [^3^H]spiperone in 40 nM ketanserin (final concentrations in assay; ketanserin added to inhibit binding to 5-HT2 receptors), then performing 2x dilutions in assay buffer (TBS containing 0.002% BSA). Assays containing tissue samples, various concentrations of [^3^H]spiperone/ketanserin, and (+)-butaclamol (2 μM; to define nonspecific binding) or assay buffer were incubated at 37°C for 1 h in a final volume of 1 ml before addition of ice-cold buffer and vacuum filtration. Thirty mice were used for radioligand binding experiments (23 males and 7 females, 17 of the **A** lineage and 13 of the **B** lineage). The density of binding sites (B_max_) and affinity of the receptors for [^3^H]spiperone (K_d_) were determined from saturation curves analysed by nonlinear regression using GraphPad Prism 9 (San Diego, California, USA). One heterozygous mouse was excluded from Figure 2 because its B_max_ value was more than 2 standard deviations from the mean. Comparisons of the heterozygous mutant group to wild type and homozygous mutant groups were both statistically significant at p < 0.05 even with that mouse.

### Stereotaxic surgery

To express GIRK2 in MSNs, *Drd2^I212F/I212F^* (9 males and 7 females) and control littermate mice (3 males and 4 females, 3-5 weeks old) were anesthetized with inhaled isoflurane (2%) and positioned in a stereotaxic apparatus. Mice were bilaterally injected with 400 nL AAV9.hSyn.tdTomato.T2A.GIRK2 (University of Pennsylvania Viral Core, V3992) into the dorsal (coordinates in millimeter from bregma: AP +1.5, ML ± 1.15, DV −4.3) and ventral (AP +1.5, ML ± 1.15, DV −4.3) part of the striatum using a Nanoject III at 100 nl/min. The pipette was kept at the site for 5 min and then slowly withdrawn. Mice were allowed to recover for at least 3 weeks following surgery.

### Midbrain Slice Preparation

Mice were anesthetized with isoflurane and euthanized by rapid decapitation. Brains were removed and placed in warm (30°C) modified Krebs buffer containing NaCl (126 mM), KCl (2.5 mM), MgCl_2_ (1.2 mM), CaCl_2_ (2.4 mM), NaH_2_PO_4_ (1.4 mM), NaHCO_3_ (25 mM), and D-glucose (11 mM) with MK-801 (3 μM). Horizontal slices containing the substantia nigra were cut at 222 μm in Krebs buffer bubbled with 95/5% O_2_/CO_2_ using a vibrating microtome (Leica). Slices were allowed to recover at 30°C in vials with 95/5% O2/CO2 Krebs with MK801 (10 μM) for at least 30 min prior to recording. Slices were hemisected and mounted in the recording chamber of an upright microscope (Olympus). The temperature was maintained at 34-36°C, and modified Krebs buffer was perfused over the slices at 1-2 ml/min. 24 mice were used in these experiments, 17 of lineage **A** and 7 of lineage **B** (11 males, 13 females).

### Slice preparation – Forebrain slices. Slice preparation – Forebrain slices

Mice were anesthetized with isoflurane and transcardially perfused with 10 mL of ice-cold carbogenated (95% v/v O2, 5% v/v CO2) sucrose cutting solution containing (in mM): 75 NaCl, 2.5 KCl, 6 MgCl_2_, 1.2 NaH2PO_4_, 25 NaHCO_3_, 0.1 CaCl_2_, 11.1 D-glucose and 1 kynurenic acid. The brain was subsequently removed and coronally sectioned (240 μM) on a vibratome. Slices were transferred to an oxygenated 34 °C chamber filled with aCSF solution consisting of (in mM): 126 NaCl, 2.5 KCl, 1.2 MgCl_2_, 2.5 CaCl_2_, 1.2 NaH_2_PO_4_, 21.4 NaHCO_3_, 11.1 D-glucose, and 10 μM MK-801 to prevent excitotoxicity for at least 1h. Afterward, slices were placed in a recording chamber which was constantly perfused with aCSF solutions at a flow rate of 2 ml/min. Solutions also contained SCH23390 (1 μM), scopolamine (200 nM), picrotoxin (100 μM), CGP55845 (300 nM), DNQX (10 μM), and dihydro-β-erythrosine hydrobromide (DHβE, 1μM). Tissue was visually identified using a BXWI51 microscope (Olympus) with custom-built infrared gradient contrast optics. Fluorescence was visualized with LEDs (Thorlabs).

### Electrophysiology – Midbrain

Recordings were obtained using glass electrodes with a starting resistance of 1.3-1.9 MΩ when filled with an internal solution containing potassium methanesulfonate (75 mM), NaCl (20 mM), MgCl_2_ (1.5 mM), HEPES potassium salt (5 mM), ATP (2 mM), GTP (0.2 mM), phosphocreatine (10 mM), and BAPTA tetrapotassium salt pH 7.35-7.45 (10 mM) at 275-288 mOsm. Cells were voltage-clamped at −60 mV using an Axopatch 200A integrating patch clamp (Molecular Devices, San Jose, CA). Recordings were made using Axograph 10 and Chart 5.5 (AD Instruments, Sydney Australia). D2 receptor-expressing dopamine neurons in the substantia nigra were identified by location, size, and firing properties. These studies were conducted by experimenters blind to mouse genotype. D2-IPSCs were elicited using a bipolar electrode and a constant current stimulus isolator (Warner Instruments, Hamden CT). CyHQ-sulpiride (Asad *et al.*, 2020) was kept as a stock solution in DMSO (10 mM) and recirculated at 15 μM. A ThorLabs M365LP1-C1 LED (Newton, NJ) was used to photolyze CyHQ-sulpiride by means of a 1 s flash (365 nm) at 6.5 mW. **Electrophysiology – Forebrain.** MSNs were voltage-clamped in whole-cell configuration at −60 mV using Axopatch 200B amplifiers (Molecular Devices, San Jose, CA), and signals were acquired with Axograph X at 5 kHz and filtered to 2 kHz or acquired with LabChart (ADInstruments; Colorado Springs, CO) at 1 kHz. Recording glass pipettes with a tip of resistance of 1.5-2 MΩ (World Precision Instruments; Sarasota, FL) were filled with potassium-based intracellular solution (in mM): 115 K-methylsulphate, 20 NaCl, 1.5 MgCl_2_, 10 HEPES(K), 10 BAPTA-tetrapotassium, 1 mg/ml ATP, 0.1 mg/ml GTP, and 1.5 mg/ml sodium phosphocreatine, pH=7.4, 275 mOsm. Dopamine release was triggered by electrical stimulation (1 ms) using a monopolar glass stimulating electrode filled with aCSF. For concentration-response curve experiments, cocaine (10 μM) was included in the recording solution to block dopamine reuptake. Dopamine was bath applied via perfusion. D2-IPSCs were evoked once per minute.

### Statistical analysis

Gait data were analyzed first by 2-way ANOVA (genotype and sex). There were no main effects of sex. A significant interaction between genotype and sex was observed for forelimb propel time (F (2, 70) = 3.24, P = 0.0449). The other measures had no significant interaction between genotype and sex, so the effect of genotype was assessed by 1-way ANOVA followed by Dunnett’s multiple comparisons test.

Statistical analyses of forebrain D2-IPSC data were performed in Prism 8 (GraphPad). Statistical significance was determined using the Mann-Whitney U test, Student’s t test, or one-way analysis of variance (ANOVA) with Tukey’s post hoc analysis.

## Acknowledgements

Support for this research was provided by the National Institute of Neurological Disorders and Stroke (R21NS117713), the National Institute on Drug Abuse (T32DA007262, R01DA004523, and R01DA35821), the National Institute for Mental Health (R21MH123085), New York University Abu Dhabi, the OHSU Shared Resources Pilot program, and the US Department of Veterans Affairs, Veterans Health Administration, Office of Research and Development, Biomedical Laboratory Research and Development (Merit Review Award BX003279). We appreciate the advice of Dr. Dineke Verbeek (University of Groningen) regarding gRNA and DNA template design and genetic analysis of putative founders, and we thank David Buck and Paul Bui for genotyping and maintenance of the two *Drd2^I212F^* lineages at the VA Portland HCS.

## SUPPLEMENTARY INFORMATION

### Supplementary File 1

Primers for genomic analysis of *Drd2^I212F^* mice

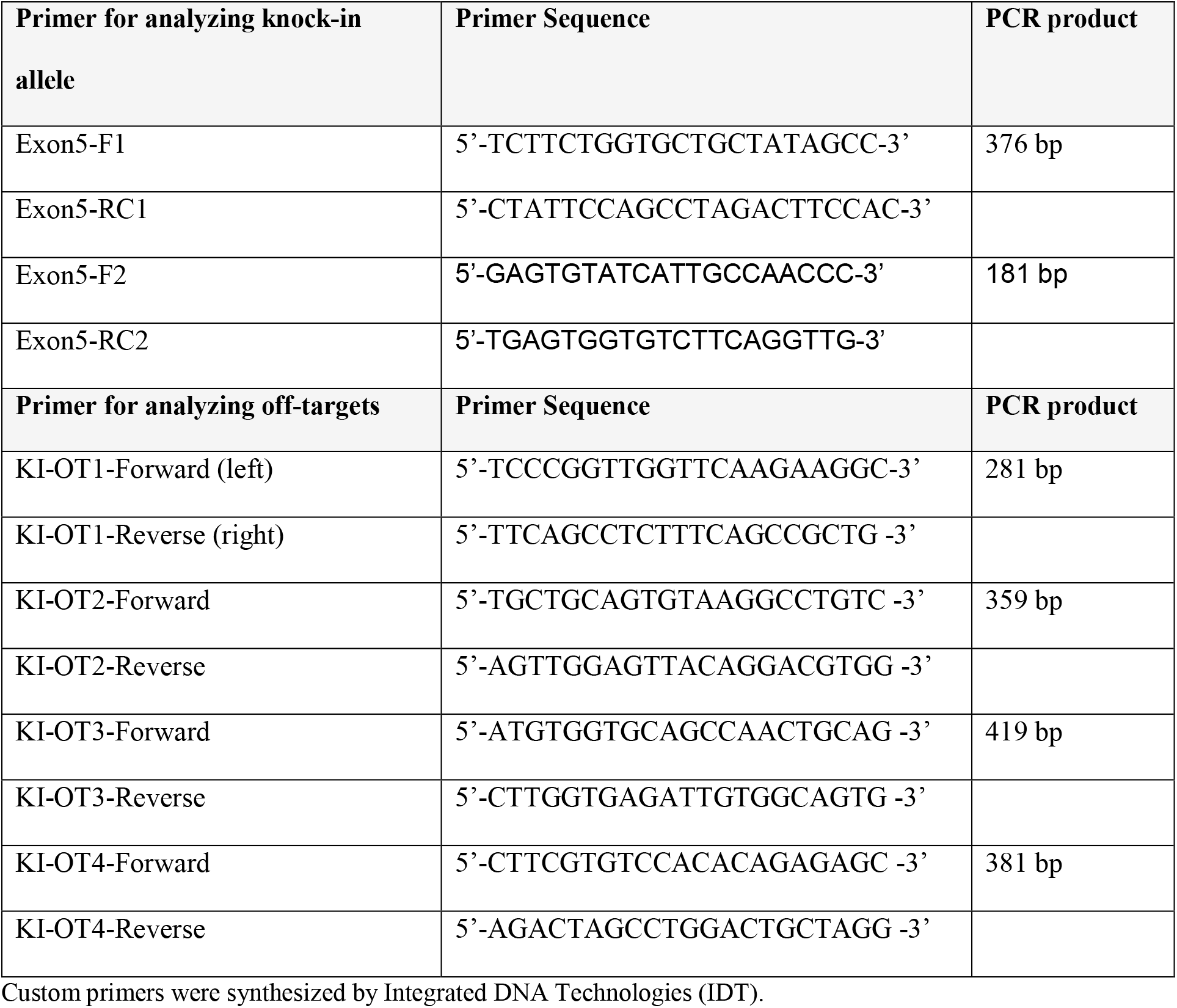

### Supplementary File 2

List of studied off-target loci for the gRNA.

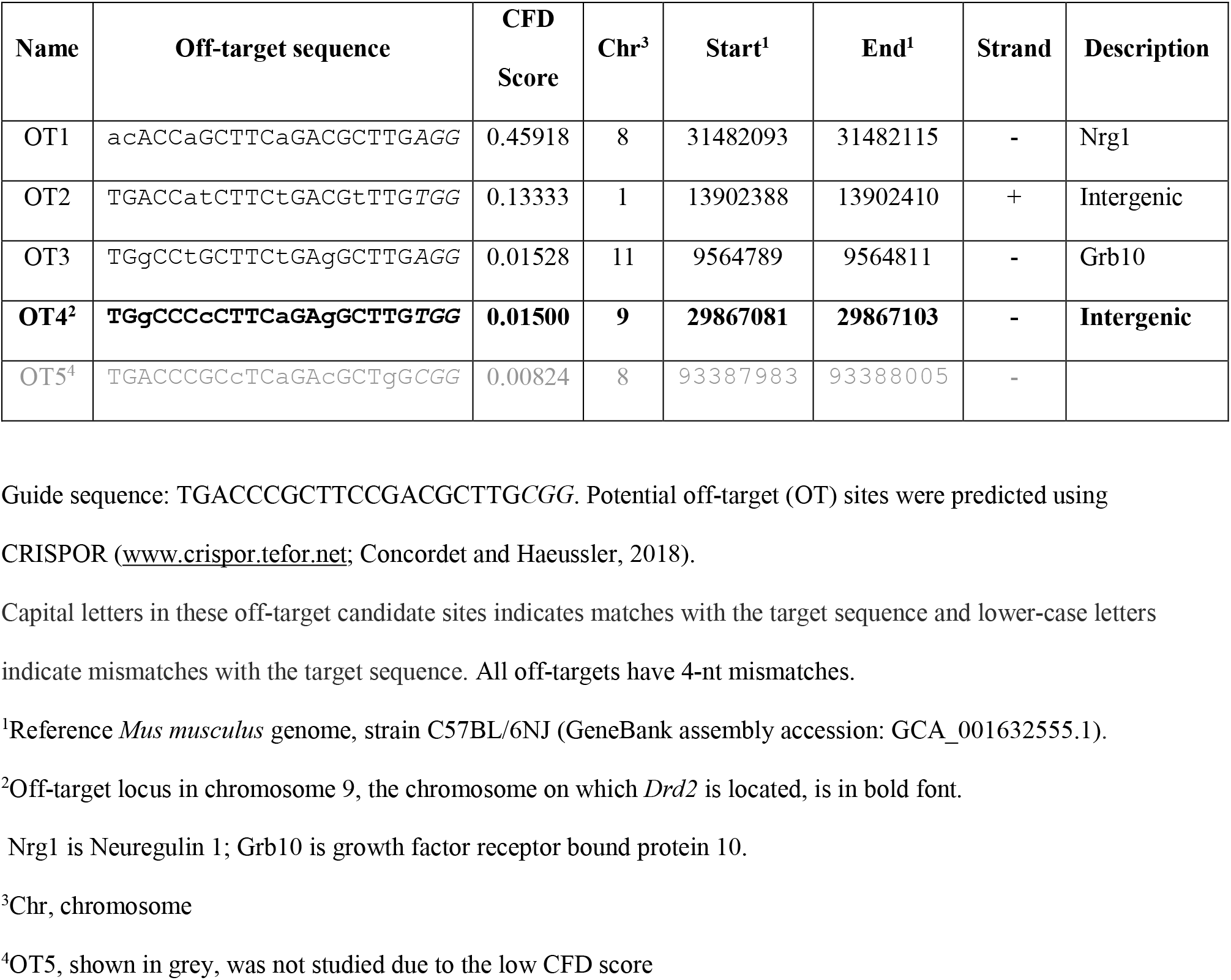

**Figure 1 – figure supplement 1.**
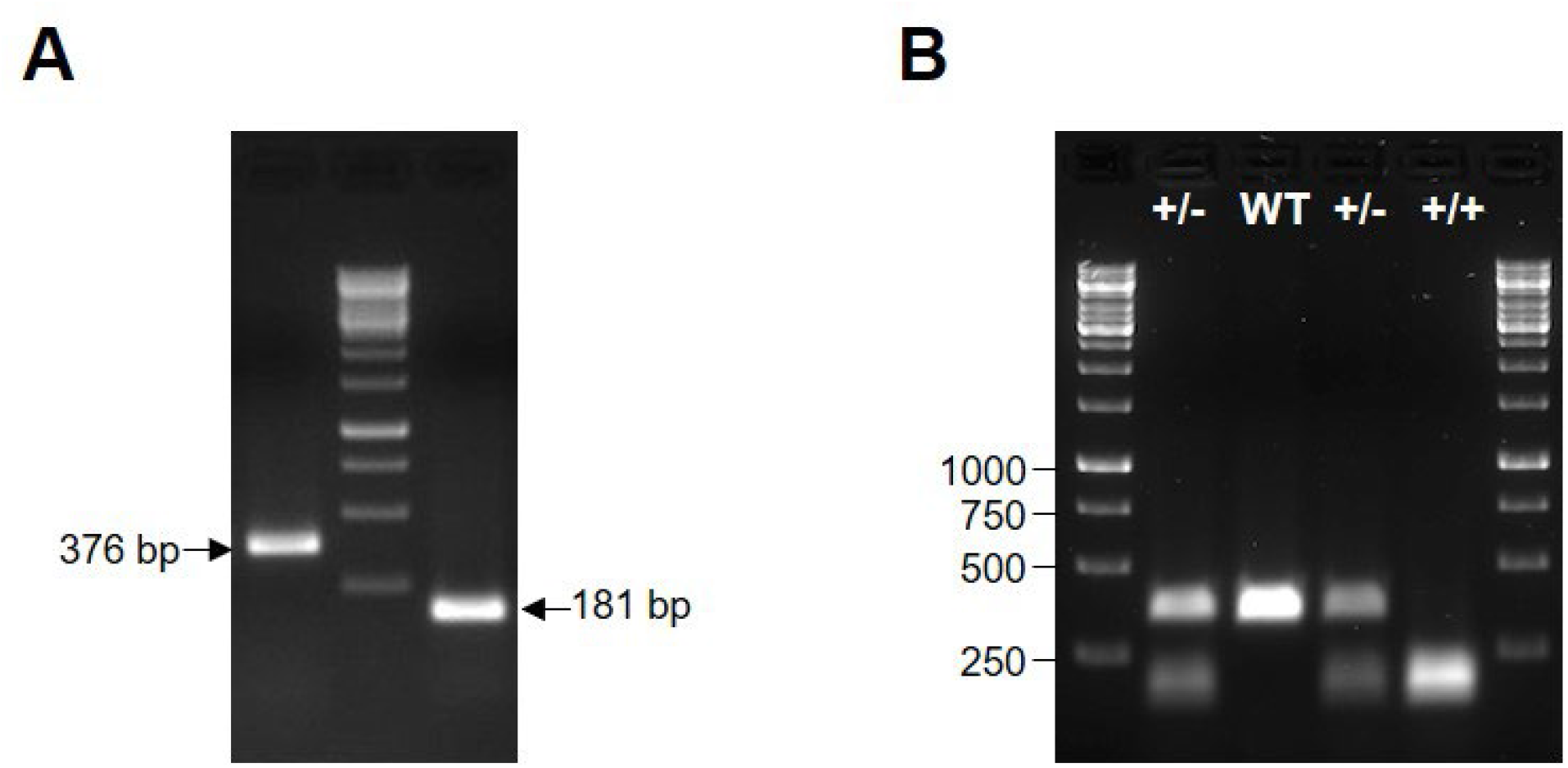
Strategy for Producing *Drd2^I212F^* mice. **A**, A 1.5% agarose gel stained with ethidium bromide showed examples of PCR products from *Drd2*-exon 5 amplified using Exon5-F1/Exon5-RC1 (376 bp, left side) and Exon5-F2/Exon5-RC2 primer pairs (181 bp, right side). **B**, A representative gel is displayed to show *Drd2^+/+^* (WT), *Drd2*^+/I212F^ (+/-) and *Drd2^I212F/I212F^* (+/+) genotypes. Representative 376-bp PCR amplicons were digested with ApoI-HF and DNA samples were run in a 1.5% agarose gel stained with SYBR Safe DNA Gel Stain (APExBIO; Houston, TX). GeneRuler 1 Kb DNA ladder (Thermo Scientific, Inc; Waltham, MA) was used as molecular weight DNA marker.

**Figure 8 – figure supplement 1.**
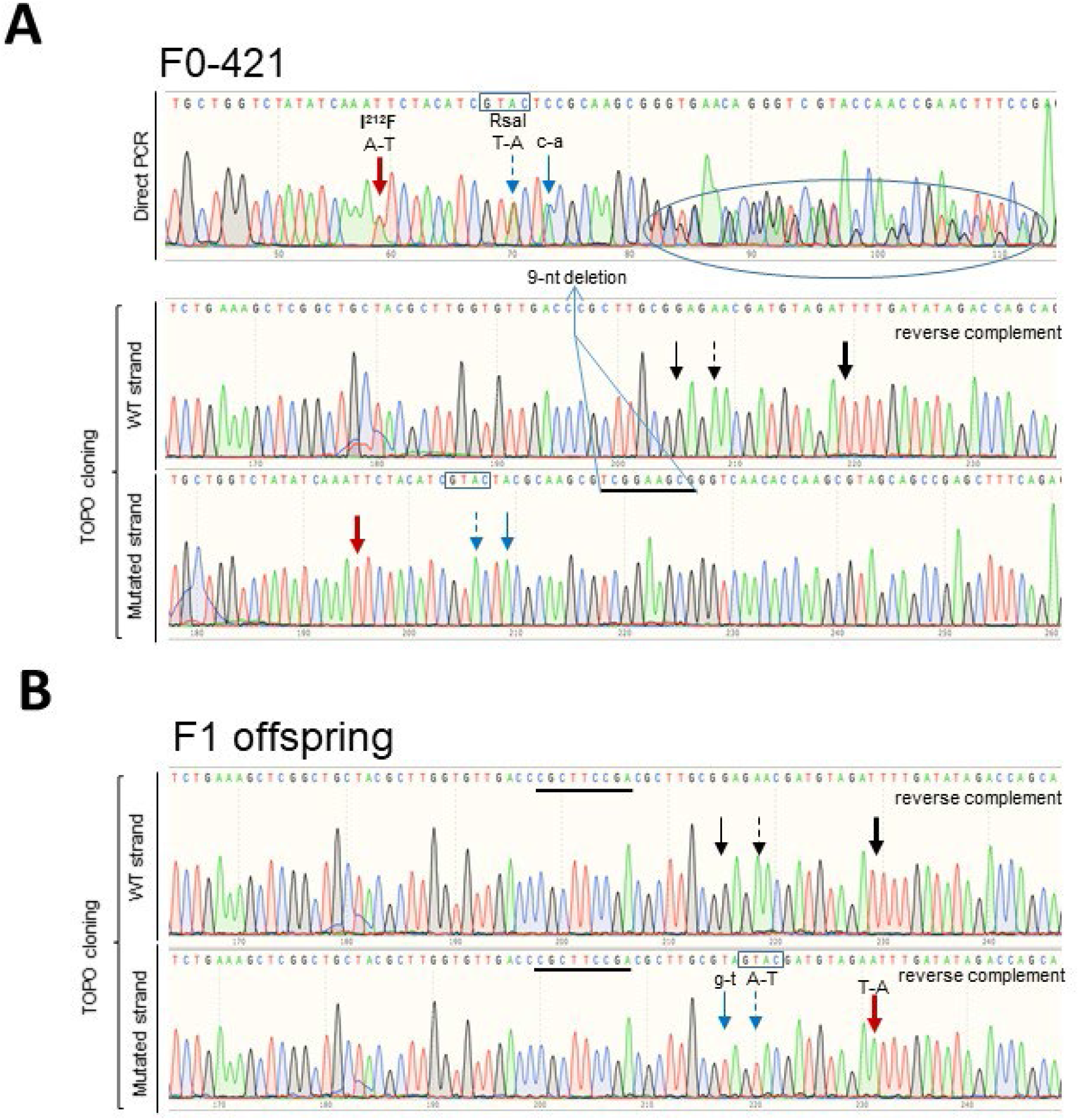
Generation of *Drd2^+/I212F^* mice: Molecular characterization of the *Drd2*-exon 5 target region in Lineage **B**. **A**, Chromatograms showing the Sanger sequencing confirmation of the c.634A>T; p.Ile212Phe in the F0-421 mouse. <u>Top</u>, the 181bp product obtained by PCR amplification of a gDNA sample using *Exon5-F2* and *Exon5-RC2* primers pair (Supplementary File 2, Figure 1) was purified and sent for sequencing. <u>Bottom</u>, 376-bp product amplified by PCR using *Exon5-F1* and *Exon5-RC1* primers pair (Figure 1) was cloned into TOPO-TA vector (Invitrogen). Plasmids containing WT and mutated strands were sent for sequencing. For the WT strand, the reverse-complemented sequence is shown. Expected nucleotide changes are present in mutated but not in WT strands; however, a deletion of nine nucleotides (underlined sequence in the mutated strand) was observed in the WT strand, explaining the mixed sequence obtained by direct sequencing of the PCR product (top chromatogram, sequence shown in an oval). **B**, Representative chromatograms showing DNA sequence of both WT and mutated strands corresponding to an F1 mouse from lineage **B**. Reverse-complemented sequences are shown. Founder F0-421 (male) was crossed with inbred control (WT) mice, and F1 offspring (eight females and eight males) were genotyped by PCR and restriction analysis. Positive F1 mice (two females and five males) were evaluated for integrity of their WT and mutated strands by cloning their 376-bp amplicon into TOPO-TA vector (Invitrogen) and sequencing both DNA strands. Wide-red arrow points to the A-T change to generate the *Drd2^I212F^* variant; blue arrows show synonymous nucleotide changes introduced downstream of the c.634A>T position. Black arrows show that no nucleotide changes are present in the WT strand. Moreover, the deleted 9-nt sequence in the founder-WT strand (A, middle chromatogram), is present in both WT and mutated strands from F1 offspring (underlined sequence, both strands). Downstream of the c.634A>T position, a specific nucleotide change (uppercase) introduced to generate an RsaI site is identified by a dashed-line arrow and the site (GTAC) is shown in a box, whereas the nucleotide change showed in lowercase was introduced to remove a Cas9 PAM sequence.

